# Antibiotic resistant bacteria survive treatment by doubling while shrinking

**DOI:** 10.1101/2024.06.27.601114

**Authors:** Adrian Campey, Remy Chait, Krasimira Tsaneva-Atanasova, Stefano Pagliara

## Abstract

Many antibiotics that are used in healthcare, farming and aquaculture end up in environments with different spatial structures that might promote heterogeneity in the emergence of antibiotic resistance. However, experimental evolution of microbes at sub-inhibitory concentrations of antibiotics has been mainly carried out at the population level which does not allow capturing heterogeneity within bacterial populations. Here we investigate and compare the emergence of resistance to ciprofloxacin in *Escherichia coli* in well mixed and structured environments using experimental evolution, genomics and microfluidics-based time-lapse microscopy. We discover that resistance to ciprofloxacin and cross-resistance to other antibiotics is stronger in the well-mixed environment due to the emergence of target mutations, whereas efflux regulator mutations emerge in the structured environment. The latter mutants also harbour sub-populations of persisters that survive high concentrations of ciprofloxacin that inhibit bacterial growth at the population level. In contrast, genetically resistant bacteria that display target mutations also survive high concentrations of ciprofloxacin that inhibit their growth via population-level antibiotic tolerance. These resistant and tolerant bacteria keep doubling while shrinking in size in the presence of ciprofloxacin and regain their original size after antibiotic removal, which constitutes a newly discovered phenotypic response. This new knowledge sheds light on the diversity of strategies employed by bacteria to survive antibiotics and poses a stepping stone for understanding the link between mutations at the population level and phenotypic single-cell responses.

## Introduction

The rise of antimicrobial resistance (AMR) poses a pressing challenge to global health with an estimated 5 million deaths associated with AMR every year globally(Murray et al. 2022). Overuse and misuse of antimicrobial drugs in human and animal health, as well as in agriculture, has accelerated the development of resistant pathogens, complicating the management of infections and threatening the foundations of modern medicine(Logan and Weinstein 2017; Dadgostar 2019; Turner et al. 2019; Dhingra et al. 2020; Cama et al. 2021). Addressing the threat of AMR requires a holistic approach, encompassing infection prevention and control measures, prudent antimicrobial use, the development of novel antibiotics, as well as understanding the evolutionary trajectories of bacteria during exposure to antibiotics(Mattar et al. 2020).

In the clinic, bacteria displaying strong antimicrobial resistance are routinely isolated, because of the obvious medical interest for such isolates(Hornischer et al. 2019). Experimental evolution of microbes instead provides complementary understanding of antimicrobial resistance by tracking the pathways that lead to the emergence of resistance, while exquisitely controlling the selection forces and physiological states at play(Remigi et al. 2019). Notably, experimental evolution demonstrated the important role of low antibiotic concentrations in the evolution of resistance. In fact, it had been traditionally assumed that only the use of antibiotic concentrations in the range between the minimum inhibitory concentration (MIC) of susceptible pathogens and the concentration that blocks growth of first-step resistant mutants led to the emergence of resistant mutants (i.e. the mutant selection window). However, recent evidence obtained via experimental evolution suggested that the mutant selection window is much wider and includes antibiotic concentrations below the MIC of the susceptible population(Gullberg et al. 2011; Jørgensen et al. 2013; Andersson and Hughes 2014; Wistrand-Yuen et al. 2018; Ching and Zaman 2020; Bawn et al. 2022).

For populations evolving in the presence of high antibiotic concentrations (i.e. strong selection pressure), survival is an immediate priority rather than fitness and therefore the options for resistance-conferring mutations are limited(Oz et al. 2014). On the other hand, at sub-inhibitory concentrations of antibiotics (i.e. weak selection pressure), bacteria can still grow and undergo a variety of evolutionary trajectories with the step-wise accumulation of resistance mutations(Wistrand-Yuen et al. 2018). In this scenario, only mutations where the fitness cost is negligible or low make the bacteria competitive, whereas resistance mutations that carry a high fitness cost are not enriched(Remigi et al. 2019). Crucially there are many environments, such as wastewater, soil, and certain body compartments where bacteria are exposed for extended periods of time to sub-inhibitory concentrations of antibiotics(Moriarty et al. 2007; Martinez 2008). In fact, a substantial fraction of antibiotics that are used to treat infections, for growth promotion in animals, aquaculture, or plant production ultimately ends up in the environment(Andersson and Hughes 2014). While many of these environments possess a spatial structure, experimental evolution of microbes at sub-inhibitory concentrations of antibiotics has been mainly carried out in well-mixed environments at the whole population level(Gullberg et al. 2011; Jørgensen et al. 2013; Wistrand-Yuen et al. 2018; Ching and Zaman 2020; Bawn et al. 2022) with few notable exceptions(Baym et al. 2016).

Understanding variations in evolutionary trajectories during exposure to sub-inhibitory concentrations of antibiotics would facilitate designing improved therapies as well as improving our understanding of resistance reservoirs in the environment. To achieve this aim, the emergence of resistance needs to be investigated both at the genotypic level (i.e. mediated by genetic mutations) and at the phenotypic level (i.e. without the emergence of mutations). In fact, recent evidence suggests a strong link between genetic resistance and persistence or tolerance, where a subpopulation or the whole population survives antibiotic exposure without the emergence of mutations(Van den Bergh et al. 2016; Levin-Reisman et al. 2017; Etthel Martha Windels et al. 2019). Moreover, there is also a need to investigate cross-resistance and collateral susceptibility resulting from the emergence of resistance to the antibiotic in use, since experimental evolution has recently allowed to discover networks of cross-resistance and collateral susceptibility during exposure to antibiotics in well-mixed environments(Jørgensen et al. 2013; Lázár et al. 2013; Oz et al. 2014; Remigi et al. 2019; Ching and Zaman 2020; Sakenova et al. 2024).

In this paper we use experimental evolution, genomics and microfluidics-based single-cell analysis to investigate the emergence of resistance to ciprofloxacin in *Escherichia coli* in both well mixed and structured environments. We chose this experimental model system because ciprofloxacin is routinely used to treat infections caused by *E. coli* in humans and animals (Kim and Hooper 2014), and ciprofloxacin resistance mechanisms are well characterised(Jørgensen et al. 2013; Oz et al. 2014; Huseby et al. 2017). In fact, ciprofloxacin targets two essential bacterial enzymes: the DNA gyrase made of two subunits, encoded in *E. coli* by *gyrA* and *gyrB*, and the topoisomerase IV made of two subunits, encoded by *parC* and *parE*(Hooper and Jacoby 2015). One of the main criticism of experimental evolution concerns the artificial nature of the experimental setups(Remigi et al. 2019). However, experimental evolution(Jørgensen et al. 2013; Oz et al. 2014; Huseby et al. 2017; Ching and Zaman 2020; Bawn et al. 2022) and clinical studies(Jalal et al. 2000; Lee et al. 2005; Maciá et al. 2006) identified similar sets of target mutations, *gyrA*, *parC* and *parE*, and efflux regulator mutations, *acrR*, *marR* and *soxR*, conferring resistance to ciprofloxacin. This diverse set of genes reflects the plasticity of bacteria for developing antibiotic resistance which limits the predictability of antibiotic resistance evolution(Palmer and Kishony 2013). Here we deepen this current understanding by simultaneously investigating cross-resistance, tolerance and persistence in mutants that emerged after exposure to sub-inhibitory concentrations of ciprofloxacin and displayed either target or off-target mutations. This new knowledge sheds light on the diversity of strategies employed by bacteria to survive environmental stressors and reveals the interplay between these different survival strategies.

## RESULTS

### Sub-inhibitory ciprofloxacin concentrations lead to the emergence of target mutations in the well-mixed environment and off-target mutations in the structured environment

We carried out evolutionary experiments on either soft agar plates (henceforth structured environment), or in shaken flasks (henceforth well-mixed environment) using, in both cases, lysogeny broth (LB) as the growth medium. Each experiment was carried out in biological triplicate by exposing *E. coli* BW25113 for 72 hours to ciprofloxacin at a concentration of either 0, 10%, 25%, 50% or 100% the minimum inhibitory concentration value measured for the *E. coli* BW25113 parental strain (MIC_PS_ = 0.015 µg ml^-1^).

We measured the bacterial growth velocity for each condition and time point and normalised these data to the bacterial growth velocity measured in the absence of ciprofloxacin at the same time point. Bacteria displayed a steady decline in growth velocity in the structured environment (Fig. S1), in contrast with a transient decline followed by full recovery in growth velocity in the well-mixed environment (Fig. S2).

We determined the level of resistance in survivor bacteria harvested at the end of each of the evolutionary experiments above by measuring the MIC fold-change compared to the parental strain. To do this, each survivor population was first grown 17h in shaken flasks containing LB and the same concentration of ciprofloxacin employed in the preceding evolutionary experiment, then sub-cultured via a 1:40 dilution in LB and the same ciprofloxacin concentration for a further 2h. Next, 8 aliquots were taken from each subculture and used in microbroth serial dilution assays (Goode, Smith, Łapińska, et al. 2021) to measure the level of resistance of each survivor population in 8 technical replicates for each biological triplicate.

We found that evolutionary experiments in both environments generated populations with resistance to ciprofloxacin that increased with the concentration of ciprofloxacin employed during the evolutionary experiments (Fig. S3). Resistance levels were significantly higher for survivor populations from the well-mixed compared with the structured environment; however, in both environments, survivor populations with distinct levels of resistance emerged both between biological replicates and within each biological replicate with resistance in range from 1 to 32-fold the MIC_PS_ value (Fig. 1).

**Figure 1.**
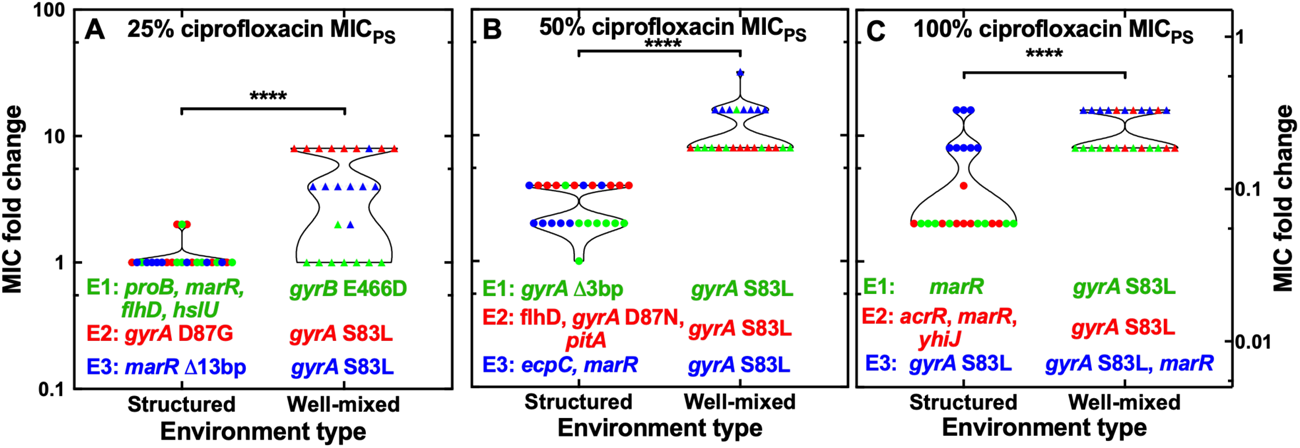
Emergence of genetic resistance to ciprofloxacin. Comparison of the emergence of resistance to ciprofloxacin in the structured (circles) and well-mixed environments (triangles) after 72 h growth in the presence of ciprofloxacin at (a) 25%, (b) 50% or (c) 100% the MIC_PS_ value. Each symbol represents the ciprofloxacin MIC value measured for one out of eight technical replicates from three different evolutionary experiments Experiment 1, 2 and 3, respectively (E1, E2 and E3, reported in green, red and blue, respectively). Corresponding mutations occurring at 100% frequency in each experiment are reported under each data set. ****: p: <0.0001.

Next, we performed whole genome sequencing on the survivor populations from each evolutionary experiment. We found that the emergence of resistance to ciprofloxacin in the well-mixed environment was underpinned by the mutation *gyrA* S83L, a single base pair substitution of C to T in the *gyrA* gene causing a serine to leucine change at amino acid position 83 of the GyrA protein (Huseby et al. 2017). This mutation was found in eight of the nine evolutionary experiments (Fig. 1 and Table S1). Interestingly, the same mutation conferred different levels of genetic resistance to ciprofloxacin in range from 4-to 16-fold the MIC_PS_ value (Fig. 1 and Table S1) and the weakest mutant further decreased its resistance in the absence of selection pressure (Fig. S4). Therefore, such heterogeneity in the level of resistance cannot be explained by differences in primary mutations but might be due to secondary mutations occurring at lower frequencies. For example, a 16-fold resistant mutant (Experiment 2 in Fig. 1C) displayed mutations in *acrR* (efflux repressor) and in *proQ* (chaperone regulator) with frequencies of 64% and 61%, respectively, besides *gyrA* S83L occurring at 100% frequency (Table S2). *gyrA* S83L was not found only in one out of nine evolutionary experiments, in which a weak resistant mutant, i.e. 2-fold the MIC_PS_ value, was underpinned by the mutation *gyrB* E466D, a single base pair substitution of A to C in the *gyrB* gene causing a glutamate to aspartate change at amino acid position 466 of the GyrB protein (Fig. 1 and Table S1). Finally, a strong resistant mutant, i.e. 16-fold the MIC_PS_ value, displayed two mutations with a single base pair deletion in the *marR* gene (efflux repressor) at nucleotide position 372, causing the removal of an adenine nucleotide, besides *gyrA* S83L (Fig. 1 and Table S1).

In striking contrast, the emergence of resistance to ciprofloxacin in the structured environment was underpinned by a much wider set of mutations (Fig. 1 and Table S1). In the first evolutionary experiment at 25% the MIC_PS_ value weak resistance to ciprofloxacin, i.e. 2-fold the MIC_PS_ value, was underpinned by the accumulation of four distinct mutations: a 11 base pair deletion in the *marR* gene at nucleotide position 241, which results in a premature stop codon; a single base pair deletion in the *proB* gene, which encodes the glutamate 5-kinase enzyme which is involved in the biosynthesis of the amino acid proline, at nucleotide position 357, causing the removal of a cytosine nucleotide; a 4 base pair insertion upstream of the *flhD* and *uspC* genes; a single base pair substitution of C to G causing a glycine to alanine change at amino acid position 208 of the HslU protein (Fig. 1 and Table S1). In the second experiment, weak resistance was conferred by the mutation *gyrA* D87G, a single base pair substitution of A to G in the *gyrA* gene causing an aspartic acid to glycine change at amino acid position 87 of the GyrA protein (Fig. 1 and Table S1). This mutant further decreased its resistance in the absence of selection pressure (Fig. S4). In the third experiment, a non-resistant mutant emerged with a 13 base pair deletion in the *marR* gene at nucleotide position 268, causing a frameshift (Fig. 1 and Table S1).

Heterogeneous genetic resistance emerged also in structured evolutionary experiments at 50% the MIC_PS_ value (Fig. 1 and Table S1). In the first experiment, weak resistance to ciprofloxacin, i.e. 2-fold the MIC_PS_ value, emerged with a 3 base pair deletion in the *gyrA* gene at nucleotide position 247, causing the removal of a serine amino acid. In the third experiment, weak resistance was conferred by two mutations: a 13 base pair deletion in the *marR* gene at nucleotide position 268, causing a premature stop codon; and a synonymous single base pair substitution of A to T in the *ecpC* gene at amino acid position 526 of the EcpC outer membrane protein (Fig. 1 and Table S1). In the second experiment, a 4-fold resistant mutant emerged with three mutations: a single base pair substitution of G to A in the *gyrA* gene causing an aspartic acid to asparagine change at amino acid position 87; a 4 base pair insertion upstream of the *flhD* and *uspC* genes; an 8 base pair deletion in the *pitA* gene at nucleotide positions 329-336, causing a frameshift (Fig. 1 and Table S1). This mutant decreased its resistance in the absence of selection pressure (Fig. S4).

Finally, there was also heterogeneity in the emergence of resistance in structured evolutionary experiments at 100% the MIC_PS_ value (Fig. 1 and Table S1). In the first experiment, a 2-fold resistant mutant emerged with a 15 base pair deletion in the *marR* gene at nucleotide position 265, causing the removal of 5 amino acids (Fig. 1 and Table S1). In the second experiment, a 4-fold resistant mutant emerged with 3 mutations: a 4 base pair insertion in the *acrR* gene at nucleotide position 217; a 20 base pair deletion in the intergenic region between *marC* and *marR*; a 20,000 base pair deletion between *yhiJ* and *yhiS* with the loss of 20 genes which have a range of roles from membrane transporter, to universal stress response and DNA damage response (Fig. 1 and Table S1). This mutant maintained its resistance in the absence of selection pressure (Fig. S4). In the third experiment, a 16-fold resistant mutant displayed the above described *gyrA* S83L mutation (Fig. 1 and Table S1).

Taken together, these data suggest that exposure to sub-MIC ciprofloxacin concentrations leads to the emergence of heterogeneous genetic resistance which is conferred by the dominant *gyrA* S83L in the well-mixed environment and by multiple target and off-target mutations including efflux pump regulators in the structured environment.

### Sub-inhibitory ciprofloxacin concentrations lead to the emergence of stronger cross-resistance in the well-mixed compared to the structured environment

Next, we set out to investigate whether the resistant mutants from the structured and well-mixed evolutionary experiments displayed genetic resistance to other antibiotics besides ciprofloxacin. We measured cross-resistance of one low and one high resistant mutant from the structured and well-mixed environments by choosing the mutant that had a ciprofloxacin MIC fold change closest to the most probable value within each dataset: the non-resistant *marR* ι113 bp mutant, the 4-fold resistant triple mutant with mutations in the genes *acrR*, *marR* and *yhiJ*, the 4-fold and 16-fold resistant *gyrA* S83L mutants, respectively (Fig. 1 and Table S1).

We found that the high resistant mutants from both the structured and well-mixed environments displayed cross-resistance to all other fluoroquinolones tested. Remarkably, the MIC values for ofloxacin (2^nd^ generation), levofloxacin (3^rd^ generation), moxifloxacin (4^th^ generation) and finafloxacin (5^th^ generation) against the high resistant mutant from the well-mixed environment were 40-fold, 40-fold, 24-fold and 64-fold higher than the corresponding MIC_PS_ values (Fig. 2a). The MIC values of these drugs against the high resistant mutant from the structured environment were also higher than the corresponding MIC_PS_ values but significantly lower than the corresponding MIC values measured against the high resistant mutant from the well-mixed environment. Surprisingly, both mutants displayed higher resistance to these fluoroquinolones compared with that measured for ciprofloxacin, that was the antibiotic used in the evolutionary experiments (Fig. 2a).

Similarly, the low resistant mutants displayed cross-resistance to the other fluoroquinolones, albeit to a lower level compared with the corresponding high resistant mutants. The MIC values for ofloxacin, levofloxacin, moxifloxacin, and finafloxacin against the low resistant mutant from the well-mixed environment were 24-fold, 8-fold, 10-fold, and 8-fold higher than the corresponding MIC_PS_ values, respectively (Fig. 2b). Interestingly, it was ofloxacin this time that showed the greatest cross resistance rather than finafloxacin which was the case for the higher resistant mutant. The MIC values of these drugs against the low resistant mutant from the structured environment were significantly lower than the corresponding MIC values measured against the high resistant mutant from the well-mixed environment (Fig. 2b).

The high resistant mutants were also resistant to representative molecules of three other antibiotic classes, beta-lactams, tetracyclines and antifolate antibiotics. Specifically, the MIC values for ampicillin, tetracycline and trimethoprim against the high resistant mutant from the structured environment were 4-fold, 4-fold and 8-fold higher than the corresponding MIC_PS_ values (Fig. 2c). The MIC values for ampicillin and tetracycline were similar against the high resistant mutants from both environments, whereas the MIC value for trimethoprim was significantly higher against the mutant from the structured environment compared with the mutant from the well-mixed environment. Interestingly, both mutants were more susceptible to representative molecules of the aminoglycosides and polymyxins classes compared with the parental strain, with the mutant from the structured environment also being more susceptible compared with the mutant from the well-mixed environment (Fig. 2c).

Similarly, the low resistant mutants displayed cross-resistance to ampicillin, tetracycline and trimethoprim and increased susceptibility to gentamicin and polymyxin B (Fig. 2d). In contrast with what observed for the high resistant mutants, this time the low resistant mutant from the well-mixed environment was more susceptible to gentamicin and polymyxin B compared with the low resistant mutant from the structured environment (Fig. 2d).

Taken together these data demonstrate that exposing bacteria to sub-inhibitory concentrations of a single antibiotic leads to the emergence of mutants that are resistant to four distinct antibiotic classes, that the level of cross-resistance is profoundly affected by the structure of the environment, and that resistance comes at a fitness cost that simultaneously renders such mutants more susceptible to two further antibiotic classes.

**Figure 2.**
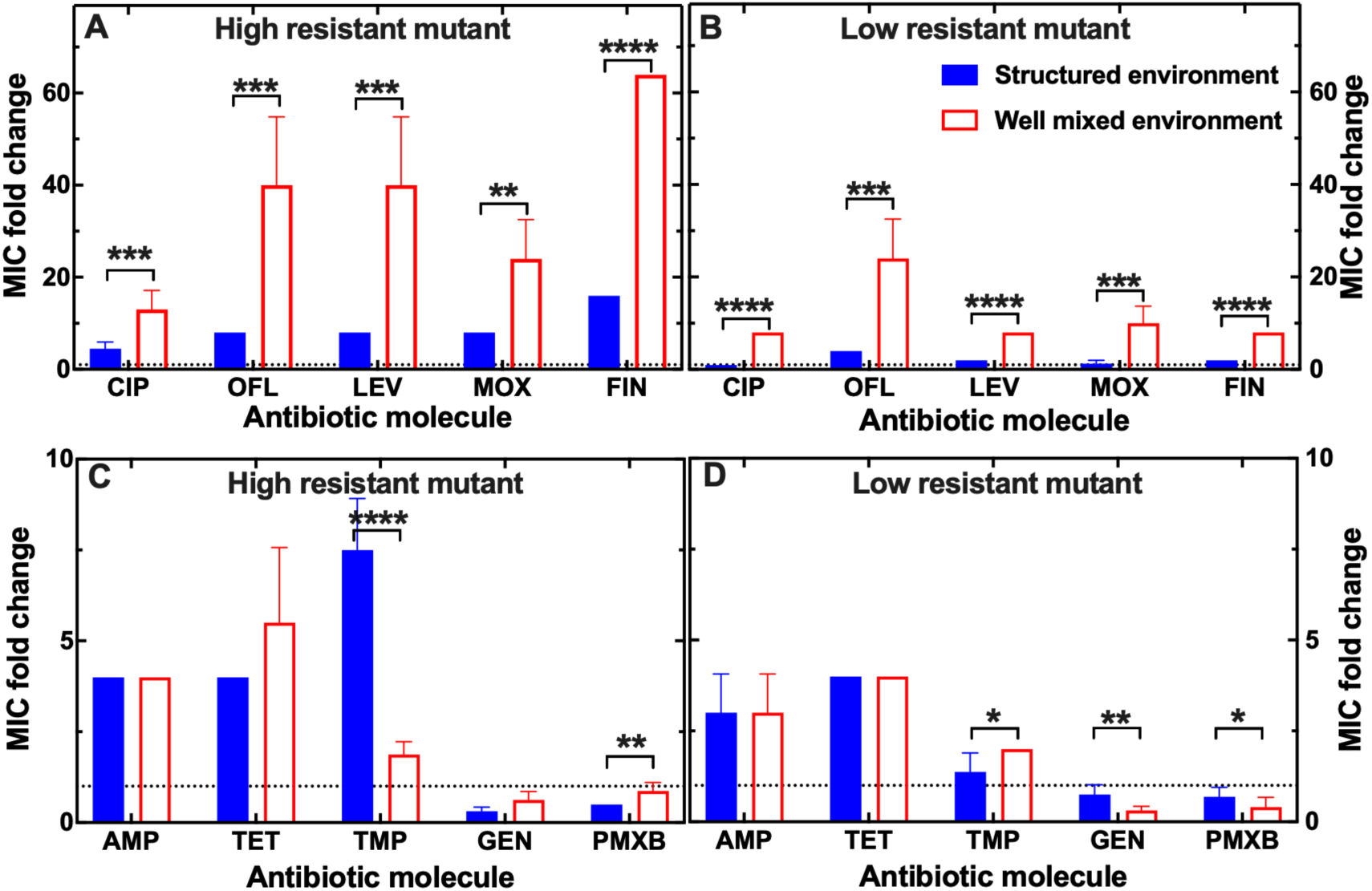
Impact of the environmental structure on the emergence of genetic multidrug resistance. Cross-resistance to other fluoroquinolones of (a) high and (b) low resistant mutants to ciprofloxacin emerged from evolutionary experiments in the structured (blue filled bars) and well-mixed environment (red empty bars). Cross-resistance to molecules from other antibiotic classes of (c) high and (d) low resistant mutants to ciprofloxacin emerged from evolutionary experiments in the structured (blue filled bars) and well-mixed environment (red empty bars). Resistance was measured as the MIC fold change compared with the MIC_PS_ and was calculated as the mean and standard error of the mean of 8 technical replicate measurements. The high and low resistant mutants were chosen as the mutants that had a ciprofloxacin MIC fold change closest to the most probable value within evolutionary experiments in the structured and well-mixed environment using ciprofloxacin at 100% and 25% the MIC_PS_, respectively. The dotted horizontal line indicates an MIC fold change of 1. *: p <0.05; **: p <0.01; ***: p <0.001; ****: p <0.0001.

### Resistant bacteria from the well-mixed environment also display a whole population tolerant phenotype

Next, we set out to determine whether besides genetic resistance to ciprofloxacin, the high resistant mutants obtained via the structured and well-mixed evolutionary experiments (i.e. the 4-fold resistant triple mutant with mutations in the genes *acrR*, *marR* and *yhiJ* and the 16-fold resistant *gyrA* S83L mutant, respectively) were also tolerant to ciprofloxacin.

As expected, we found that the mutant from the structured evolutionary experiment grew unaffected by the presence of ciprofloxacin at the MIC_PS_ value (Fig. 3a). In contrast, exposing this mutant to ciprofloxacin at 10×, 25× or 100× the MIC_PS_ value (i.e. approx. 2×, 6× or 25× the MIC_S_ value) killed the majority of the mutant population within 5 h but revealed the typical biphasic killing kinetics and the presence of persisters that constituted 0.3%, 0.06% or 0.0003% of the mutant population, respectively (Fig. 3a).

As expected, the mutant population from the well-mixed evolutionary experiment was not affected by exposure to ciprofloxacin at 1× or 25× the MIC_PS_, i.e. approx. 0.06× or 2× the MIC_WM_ (Fig. 3b). In contrast, exposing this mutant to ciprofloxacin at 100× or 200× the MIC_PS_ value (i.e. approx. 6× or 12× the MIC_WM_ value) gradually killed the mutant population (Fig. 3b). These data suggest that the whole bacterial population was tolerant to ciprofloxacin treatment rather than containing a sub-population of persisters, as it was the case for the mutant from the structured environment evolutionary experiment (Fig. 3a).

Taken together these data suggest that, besides genetic resistance to ciprofloxacin, the mutant obtained from the structured evolutionary experiment contains subpopulations of persisters to ciprofloxacin, whereas the mutant obtained from well-mixed evolutionary experiment displays tolerance to ciprofloxacin at the population level.

**Figure 3.**
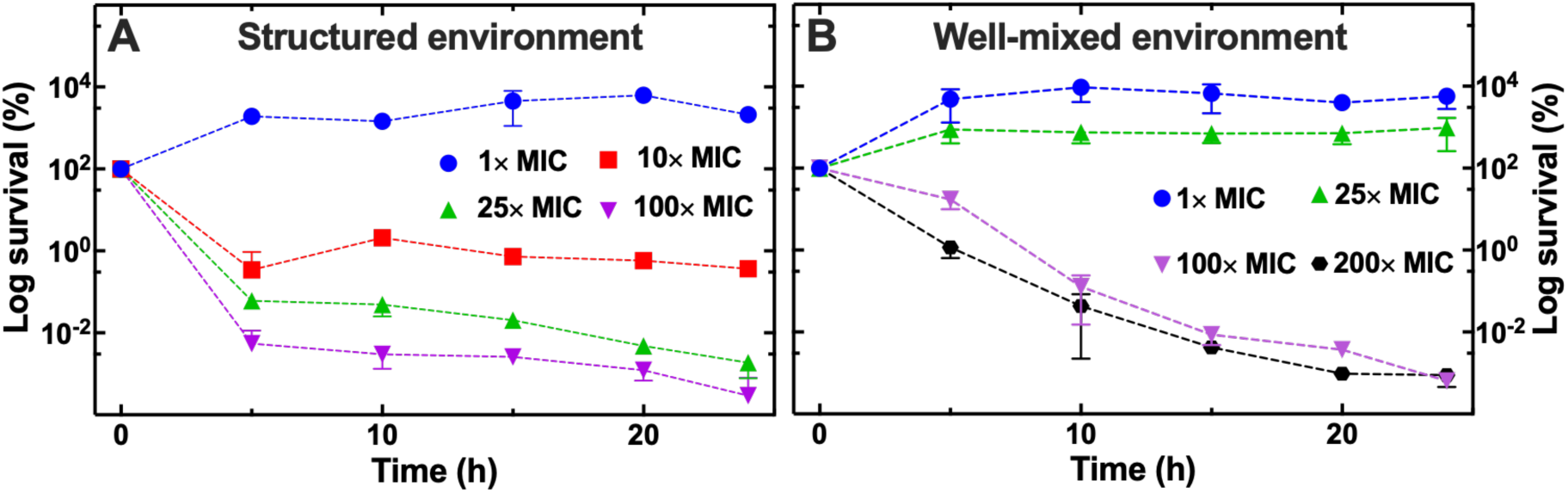
Impact of the structure of the environment on tolerance to ciprofloxacin within genetically resistant populations. Temporal dependence of bacterial counts for the high resistant mutants obtained from evolutionary experiments using 100% the MIC_PS_ value in (a) the structured and (b) the well-mixed environment after exposure to 1× (blue circles), 10× (red squares), 25× (green upward triangles), 100× (purple downward triangles) or 200× (black circles) the MIC_PS_ value. The log survival at each time point was calculated as the mean and standard deviation of the ratio of the colony forming units at that time point divided by the corresponding colony forming units at t = 0 performed in biological triplicate.

### Resistant bacteria from the well-mixed environment survive treatment by doubling while shrinking

Next, we set out to investigate the phenotype of the high resistant mutants from both environments (i.e. the 4-fold resistant triple mutant with mutations in the genes *acrR*, *marR* and *yhiJ* and the 16-fold resistant *gyrA* S83L mutant) in terms of single-cell growth before, during and after exposure to ciprofloxacin. We employed our recently introduced high-throughput, microfluidics-based platform to perform kinetic analysis of antibiotic efficacy (Bamford et al. 2017; Goode, Smith, Zarkan, et al. 2021; Goode, Smith, Łapińska, et al. 2021; Cama et al. 2022; Zhang et al. 2023) and accumulation (Blaskovich et al. 2019; Cama et al. 2020; Stone et al. 2020; Łapińska et al. 2022) in individual bacterial cells. Briefly, we used a microfluidic mother machine device equipped with thousands of compartments physically separated from each other, each compartment initially hosting one bacterium from an overnight culture of either the structured or well-mixed high resistant mutant. We supplied LB medium via microfluidics to the environment around the compartments for 120 min and imaged single-cell growth of both mutants (Fig. 4a-b). We found that during this period the two mutants displayed similar bacterial lengths and doubling times (Fig. 4c-d and 4g-h), whereas the mutant from the well-mixed environment displayed a significantly faster elongation rate (p < 0.01, Fig. 4e-f).

Next, we supplied LB medium containing ciprofloxacin at 25× the MIC_PS_ value for 240 min (i.e. around 6× and 2× the MIC value measured for the structured and well-mixed resistant mutant, respectively). During this period the average bacterial length of the mutant from the structured environment first significantly increased from 3.8 µm at t = 120 min to 6.0 µm at t = 240 min (p < 0.0001) and then slightly decreased to 5.4 µm at t = 360 min (Fig. 4c). In stark contrast, the average bacterial length of the mutant from the well-mixed environment significantly decreased from 3.8 µm at t = 120 min to 3.1 µm at t = 240 min and 2.5 µm at t = 360 min (p < 0.0001, Fig. 4d). The two mutants also displayed a strikingly different response to ciprofloxacin treatment in terms of elongation rate. The average elongation rate significantly decreased for the mutant from the structured environment from 2.3 µm h^-1^ to 0.6 µm h^-1^ and 0.2 µm h^-1^ (at t = 120, 240 and 360 min, respectively, p < 0.0001, Fig. 4e). In contrast, the average elongation rate for the mutant from the well-mixed environment firstly significantly decreased from 3.1 µm h^-1^ to 0.3 µm h^-1^ and then increased to 1.2 µm h^-1^ (at t = 120, 240 and 360 min, respectively, p < 0.0001, Fig. 4f). Finally, the average doubling time of the mutant from the structured environment significantly increased from 105 min to 180 min and 307 min (at t = 120, 240 and 360 min, respectively, p < 0.0001, Fig. 4g). In contrast, the average doubling time of the mutant from the well-mixed environment first significantly decreased from 114 min to 90 min and then significantly increased to 148 min (at t = 120, 240 and 360 min, respectively, p < 0.0001, Fig. 4h). Therefore, the two mutants displayed two distinct responses to exposure to supra-MIC concentrations of ciprofloxacin: cells of the mutant from the structured environment became longer while elongating and doubling slowly; cells of the mutant from the well-mixed environment became shorter while elongating and doubling fast.

Finally, we supplied again LB medium for 240 min, thus removing extracellular ciprofloxacin from the microfluidic environment. During this period the average bacterial length of the mutant from the structured environment did not significantly change (Fig. 4c), the average elongation rate significantly decreased down to 0.1 µm h^-1^ (p < 0.05, Fig. 4e) and none of the bacteria doubled (Fig. 4g). In contrast, both the average bacterial length and elongation rate of the mutant from the well-mixed environment significantly increased up to 11.4 µm and 10.5 µm h^-1^ at t = 600 min (p < 0.0001, Fig. 4d and 4f), whereas its doubling time firstly increased from 148 min to 214 min and then decreased down to 134 min (at t = 360, 480 and 600 min, respectively, p < 0.0001, Fig. 4h). Therefore, the two mutants displayed two distinct responses after exposure to supra-MIC concentrations of ciprofloxacin: cells of the mutant from the structured environment stopped growing; cells of the mutant from the well-mixed environment became longer while continuing to elongate and double. Noteworthy, the mutant from the well-mixed environment displayed large growth heterogeneity: some cells became very long displaying a filamenting phenotype and divided very slowly (Bos et al. 2015; Goormaghtigh and Van Melderen 2019; Etthel M. Windels et al. 2019), whereas other cells displayed length, elongation rate and doubling time comparable to those measured before exposure to ciprofloxacin (Fig. 4).

**Figure 4.**
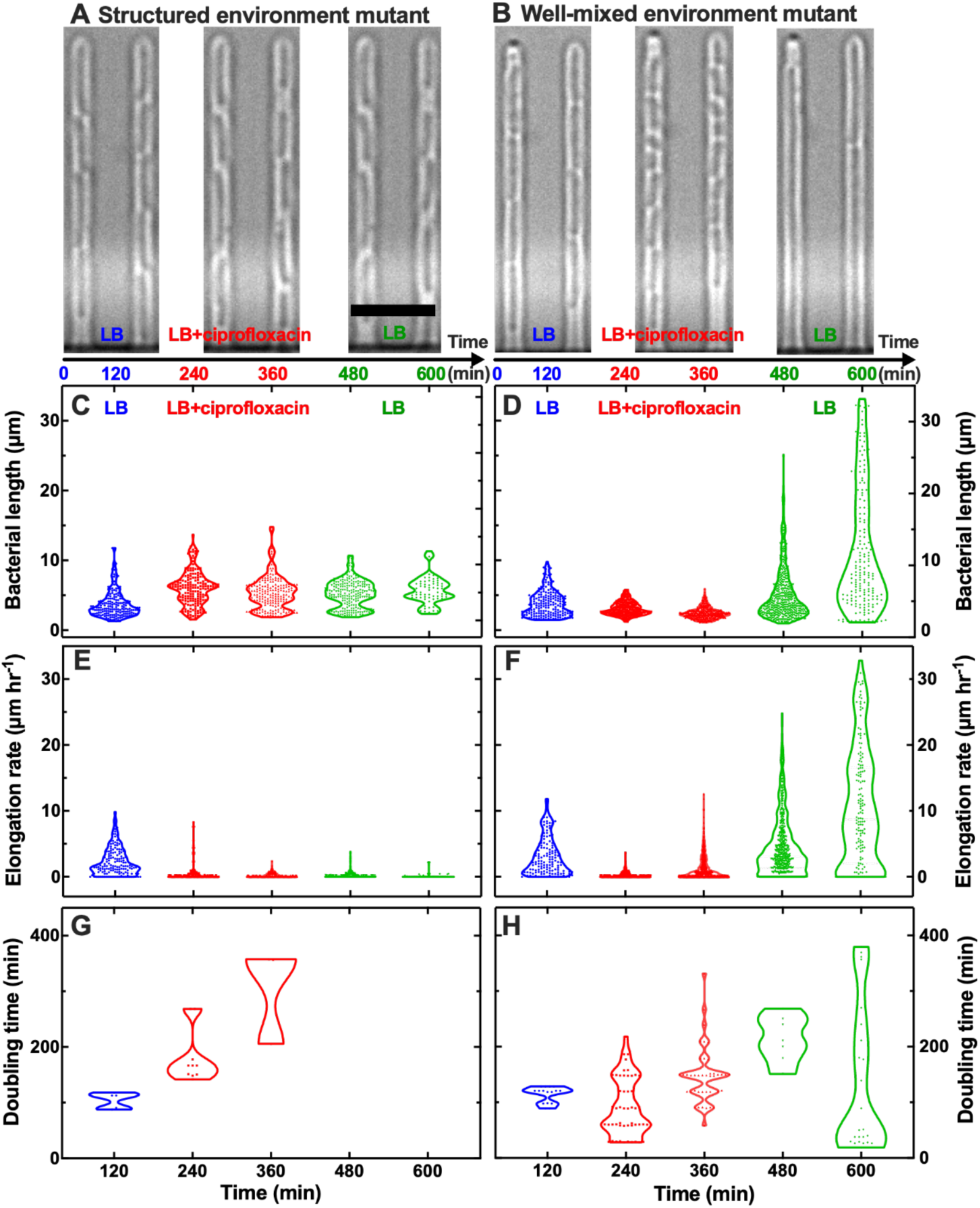
Resistant mutants from the well-mixed environment survive ciprofloxacin treatment by shrinking while doubling. Representative microscopy images of the high resistant mutant from (a) the structured environment and (b) the well-mixed environment during exposure to LB growth medium, ciprofloxacin at a concentration 25× the MIC_PS_ value, and again LB growth medium. Scale bar: 5 µm. Corresponding (c-d) bacterial length, (e-f) elongation rate and (g-h) doubling time for individual bacteria from the high resistant mutant populations from the structured environment and the well-mixed environment during exposure to LB growth medium (0 < t < 120 min, blue dots), ciprofloxacin at a concentration 25× the MIC_PS_ value (120 < t < 360 min, red dots) and again to LB growth medium (360 < t < 600 min, green dots). Each dot represents a measurement carried out on an individual bacterium. We carried out these measurements starting from 15 individual bacteria from each mutant from biological triplicate at t = 0 and took regular measurements on these bacteria and their progeny every 10 min. For clarity, we collated these measurements in violin plots at 120 min intervals.

## Discussion

### Impact of the environmental structure on bacterial resistance

Experimental evolution of microbes at sub-inhibitory concentrations of antibiotics in well-mixed environments has recently demonstrated the important role of low antibiotic concentrations in the evolution of resistance (Gullberg et al. 2011; Jørgensen et al. 2013; Wistrand-Yuen et al. 2018; Ching and Zaman 2020; Bawn et al. 2022). However, many natural environments possess a spatial structure. Using experimental evolution of *E. coli* at sub-inhibitory concentrations of ciprofloxacin, we found a threshold concentration for resistance development at 25% MIC_PS_ value. This is in contrast to previous research that has reported the minimum selective concentration of ciprofloxacin at 10% the MIC_PS_ value or below, albeit in different experimental conditions with longer exposure (Gullberg et al. 2011; Stanton et al. 2020). Above this threshold concentration value, resistance increased with the antibiotic concentration employed in the evolutionary experiments. This observation aligns with the concept of a concentration-dependent response, where the bacterial response varies with the intensity of the antibiotic stress (Bernier et al. 2013; Baquero and Levin 2021).

A particularly striking observation was the differential levels of antibiotic resistance development between the two environmental conditions tested. *E. coli* bacteria in the well-mixed environment consistently showed higher average levels of resistance compared to their counterparts in the structured environment. The increased levels of antibiotic resistance in the well-mixed environment could be attributed to a range of factors, such as more uniform exposure to antibiotics or quicker dissemination of resistant mutants or resistant genes through horizontal gene transfer (Cerca et al. 2005). This finding will help understanding of how antibiotic resistance may develop differently in various bodily or natural environments, since the well-mixed environment broadly recapitulates bloodstream or urinary tract infections and aquatic natural environments (van der Poll et al. 2021), whereas the structured environment broadly recapitulates biofilm-associated infections on indwelling medical devices or in the lungs and soil (Turcios 2020).

Notably, compared to previous studies (Jørgensen et al. 2013; Oz et al. 2014; Wistrand-Yuen et al. 2018), we observed a more rapid onset of antibiotic resistance under sub-lethal antibiotic concentrations across both environments, implying that even short-term exposure to antibiotics can significantly hasten the evolution of resistant bacterial strains.

### Emergence of heterogeneous genetic resistance in structured environments

The target of ciprofloxacin is the DNA gyrase, which is formed of two subunits encoded by *gyrA* and *gyrB* in *E. coli*, and is responsible for cleaving and re-joining the DNA strands during the process of supercoiling (Cozzarelli 1980). Mutations in *gyrA*, particularly *gyrA* S83L are significantly overrepresented among both fluoroquinolone resistant clinical isolates and resistant mutants generated via experimental evolution (Oz et al. 2014; Huseby et al. 2017). Accordingly, we found the *gyrA* S83L mutation in eight out of the nine mutants from the well-mixed environment investigated. Interestingly, this same mutation conferred different levels of genetic resistance to ciprofloxacin suggesting that other mechanisms might be involved in the emergence of resistance. In striking contrast only one out of the nine mutants from the structured environment investigated carried the *gyrA* S83L mutation. These findings are remarkable as they demonstrate the crucial impact of the structure of the environment on the emergence of genetic resistance to antibiotics, alongside the well-established impact on resistance to phage (Brockhurst et al. 2004; Gómez and Buckling 2011; Hernandez and Koskella 2019; Attrill et al. 2021; Attrill et al. 2023).

Exposure to sub-inhibitory concentrations of ciprofloxacin in the structured environment led to the emergence of heterogeneous genetic resistance highlighting the importance of investigating sub-populations within bacterial cultures for emerging resistance traits. Five mutants carried off-pathway mutations in the *marR* gene. *MarR* encodes the transcriptional repressor of the transcription factor MarA, which in turn controls several genes involved in resistance to antibiotics including the AcrAB–TolC efflux pump (Holden et al. 2023). Only one of these mutants additionally carried a mutation in *acrR*, encoding the repressor AcrR that negatively regulates *acrAB* (Holden et al. 2023), suggesting that acrR might play a lesser role compared to *marR* in terms of resistance to ciprofloxacin in the structured environment. Four mutants carried target mutations, such as the above mentioned *gyrA* S83L that conferred the highest level of resistance to ciprofloxacin recorded in the structured environment, or *gyrA* D87N and D87G mutations that were previously found to be additional mutations in some *gyrA* S83L mutants (Huseby et al. 2017). Some mutants carried additional off-pathway mutations in genes that are not thought to be involved in antibiotic resistance such as *proB*, encoding the glutamate 5-kinase that facilitates proline biosynthesis, *hlsU* which encodes the ATPase component of the HslVU protease, *flhD* encoding a master regulator of several flagellar genes, or *uspC* involved in the universal stress response. It is conceivable that some of these mutations might enhance bacterial fitness in the physically constrained structured environment (Nachin et al. 2005), further highlighting the role of population heterogeneity in conferring antibiotic resistance (Baquero and Levin 2021). Finally, we did not detect the mutation *parC* S80I in any of the mutants from either environments, despite this mutation being most frequently selected after the initial selection of *gyrA* S83L in a previous experimental evolution study (Huseby et al. 2017), suggesting that *parC* S80I emerges predominantly during exposure to high fluoroquinolone concentrations.

Taken together these findings demonstrate that multiple evolutionary trajectories can lead to genetic resistance to antibiotics and question diagnostic methods that exclusively rely on detecting specific high-frequency mutations to infer resistance levels.

### Impact of the environmental structure on the emergence of cross-resistance

High levels of cross resistance are known to occur within the fluoroquinolone antibiotic class (Batard et al. 2016; Ching and Zaman 2020) and our results show that this alarming phenomenon also emerges after sub-inhibitory antibiotic exposures and across different environments. The observed cross-resistance to finafloxacin is of particular concern, considering that this new fluoroquinolone was designed to overcome infections that are resistant to older fluoroquinolones (Kocsis et al. 2021), thus challenging the assumption that newer antibiotics can outpace bacterial adaptation (Gray and Wenzel 2020).

In accordance with our resistance data, the *gyrA* S83L mutant emerging from the well-mixed evolutionary experiments also displayed stronger cross-resistance to other fluoroquinolones compared to the triple mutant with mutations in the genes *acrR*, *marR* and *yhiJ* emerging from the structured evolutionary experiments. These mutants also displayed similar cross-resistance levels to ampicillin and tetracycline (from the beta-lactam and tetracycline class, respectively), whereas the mutant from the structured environment displayed higher levels of cross-resistance to trimethoprim (from the antifolate class). These data therefore suggest that the mechanisms of cross resistance against other antibiotic classes are not dependent on the type and level of resistance to ciprofloxacin but may be part of a more general bacterial response to the exposure to ciprofloxacin (Mathieu et al. 2016), and might complicate efforts to rotate antibiotics as a strategy to mitigate resistance (Chatzopoulou and Reynolds 2022).

Finally, we observed collateral sensitivity to gentamicin (from the aminoglycoside class) and polymyxin B (from the polymyxin class) in the mutants from both environments, in accordance with previous studies suggesting that some antibiotics might become more effective as second-line treatments or in combination therapies (Chait et al. 2007; Pál et al. 2015).

### Phenotypic resistance within genetically resistant bacterial populations

Antibiotic tolerance and persistence are generally investigated within genetically susceptible bacterial populations (Bamford et al. 2017; Balaban et al. 2019) with recent evidence suggesting that these surviving bacteria constitute a pool for the emergence of genetic resistance (Levin-Reisman et al. 2017; Barrett et al. 2019; Etthel M Windels et al. 2019; Brandis et al. 2023). Here we complement these recent efforts by demonstrating that both antibiotic tolerance and persistence play a key role also within genetically resistant populations and are not a key feature only in genetically susceptible populations. Therefore, these new data demonstrate that bacterial populations simultaneously use both genetic and non-heritable resistance to overcome exposure to antibiotics.

Notably, the structure of the environment drives a switch between antibiotic tolerance at the population level and persistence at the sub-population level. The typical biphasic killing kinetics observed in the triple mutant with mutations in the genes *acrR*, *marR* and *yhiJ* emerging from the structured evolutionary experiments reveals the presence of a small subpopulation of persister cells (Brauner et al. 2016). These cells are capable of surviving antibiotic concentrations that are lethal to the majority of the bacterial population. This finding aligns with existing literature that recognises persisters as phenotypic variants that make up a small percentage of bacterial populations (Balaban et al. 2019) and contribute to the recalcitrance of infections, particularly in structured environments like biofilms (Wainwright et al. 2021). It is conceivable that stress responses induced by exposure to ciprofloxacin, such as the SOS response, are involved in the development of both persistence and genetic resistance (Soares et al. 2020; Crane et al. 2021). Accordingly, we found that resistant mutants from the structured environment displayed mutations of transcriptional regulators *marR* and *acrR* involved in stress responses and efflux that have been shown to play a key role in persistence (Pu et al. 2016; Łapińska et al. 2022; Lee et al. 2024).

In contrast, the *gyrA* S83L mutant emerging from the well-mixed evolutionary experiments displayed a remarkable population level tolerance to ciprofloxacin, with a slow, plateaued killing at higher concentrations. This slower, steady killing, suggests a population-level adaptation at both the genetic and phenotypic level that allows a larger fraction of cells to withstand exposure to high concentrations of antibiotics (Westblade et al. 2020). We discovered that these genetically resistant and antibiotic tolerant bacteria survive high concentrations of ciprofloxacin by displaying an unusual phenotypic response: immediate arrest in bacterial elongation with simultaneous continued cell division leading to shrinking of bacterial cells. This newly observed response may represent an adaptive strategy, where the bacteria prepare for the possibility of a less hostile environment in the future. Recent evidence suggested that antibiotics induce distinct morphological changes depending on their cellular targets, DNA targeting antibiotics, such as ciprofloxacin, leading to filamentation and an increase in cell length (Goormaghtigh and Van Melderen 2019; Cylke et al. 2022). However, our data clearly show a contrasting result, highlighting the need for further research on the impact of resistance to fluoroquinolones on the cell cycle (Jonas 2014) considering that mutations in *gyrA* are significantly overrepresented among fluoroquinolone resistant clinical isolates (Oz et al. 2014; Huseby et al. 2017).

In conclusion, our data reveal the diversity of strategies employed by bacteria to survive antibiotics and demonstrate how the structure of the environment influences the interplay between genetic and phenotypic resistance mechanisms with implications for the future development of effective antibiotic treatment strategies.

## Materials and Methods

### Chemicals and bacterial culture

All chemicals were purchased from Fisher Scientific or Sigma-Aldrich unless otherwise stated. Bacteria were cultured in lysogeny broth (LB) medium (10 g/L tryptone, 5 g/L yeast extract, and 0.5 g/L NaCl; Formedium). LB agar plates with 15 g/L agar were used for streak plates. Stock solutions of ciprofloxacin were prepared by dissolving it in 0.1 M HCL in Milli-Q water and serially diluting to experimental concentrations in LB on the day of the experiment. The parental *Escherichia coli* strain BW25113 was purchased from Dharmacon (GE Healthcare) and stored in 50% glycerol stock at -80 °C. Streak plates for this strain were made by thawing a small aliquot of the glycerol stock and plated onto LB agar plates. Overnight cultures of *E. coli* BW25113 were prepared by inoculating a single bacterial colony from a streak plate in a glass flask containing 100 ml of LB and incubating it on a shaking incubator at 200 rpm at 37 °C for 17 hours.

Culturing of previously exposed resistant mutant strains was performed in the same manner except for the addition of ciprofloxacin into the LB medium, at the same concentration that was used in the generation of the mutants, in order to maintain the same selection pressure.

### Evolutionary experiments in the well-mixed environment

An overnight culture of *E. coli* BW25113 was diluted 1:100 into 100 mL of fresh LB which contained ciprofloxacin at a concentration of either 0.015 μg mL^-1^, 0.0075 μg mL^-1^, 0.00375 μg mL^-1^, 0.0015 μg mL^-1^, or no ciprofloxacin to give 100%, 50%, 25%, 10%, and 0% of the parental strain MIC value (MIC_PS_), respectively. Cultures were then grown for 24 hours on a shaking incubator at 200 rpm at 37 °C. After this, the cultures were diluted 1:100 daily into 100 mL of fresh medium containing the same antibiotic concentration for a total time of 72 hours. At this point three 1 mL aliquots of each culture were stored in 50% glycerol at -80 °C. Throughout each 72 hour long evolutionary experiment, samples were taken at regular time points and OD_600_ measurements performed to measure the growth velocity of each *E. coli* culture under each ciprofloxacin exposure.

### Evolutionary experiments in the structured environment

“Swim” agar plates were made by pouring 0.3% LB agar into petri dishes with a 1-50 numbered grid sticker on them (Merck). Ciprofloxacin was added from a stock solution to each plate to a final concentration of either 0.015 μg mL^-1^, 0.0075 μg mL^-1^, 0.00375 μg mL^-1^, 0.0015 μg mL^-1^, or no drug to give 100%, 50%, 25%, 10%, and 0% MIC_PS_, respectively. Each plate was inoculated with a single colony of *E. coli* BW25113 from a streak plate at two sites, using the numbered grid background to ensure consistency of inoculation site across the plates. The plates were then loaded onto a plate scanner device (Epson Perfection V800 Colour Image Scanner, used in transparency mode) and housed in a custom-built incubator set to 37 °C. Time lapse images were acquired by a Raspberry Pi computer attached to the scanner running a custom python script that used the SANE imaging package. Images were captured every 5 minutes over a period of 72 hours in order to visually monitor bacterial growth dynamics. Images were processed using ImageJ software to analyse both area growth and velocity of the chemotactic front. Firstly, images were imported as a stack and the scale of the images was worked out using the known distance of the numbered gridded squares on swim agar plates. To measure the area growth, the original inoculation site at t = 0 was outlined with the free hand drawing tool and measured to give the area in mm^2^. This step was repeated at the end of the experiment (t = 72 hours), along the whole edge of the total area of growth. Fold increase was calculated by dividing the end point (t = 72 hours) area growth by the start point (t = 0) area growth. In order to measure the velocity of the chemotactic front, images were split into respective red, green, and blue image components. The red split was chosen to conduct further analysis as it showed the highest visual contrast. The brightness and contrast of the images was then adjusted to further enhance the contrast in order to see the position of the front and these settings were applied to all images in the stack. Next, a region of interest (ROI) line was marked, running perpendicular to the front and in the direction of travel and the multikymograph tool was used to generate a kymograph. The gradient of this kymograph was marked with another ROI and then the velocity measurement tool was used to generate a measurement for the mean velocity for each segment in mm hr^-1^. This data was then normalised by dividing each measurement by the measurement for the no drug control at each time point. This data was then analysed and plotted using GraphPad prism 9.

At the end of each evolutionary experiment, samples from each plate were inoculated in 100mL fresh LB medium containing the same ciprofloxacin concentration used throughout the evolutionary experiment. 1 mL aliquots from each overnight culture were used to produce 50% glycerol stocks that were stored at -80 °C.

### Measurement of the minimum inhibitory concentration

Minimum inhibitory concentration (MIC) assays were determined using the broth microdilution method (Smith et al. 2018). First, a 17-hour overnight culture was diluted 1:40 in Mueller Hinton broth (MHB), adding ciprofloxacin at the appropriate concentration when necessary, and grown to exponential phase (OD_600_ = 0.5). Next, 100 μL of 2 μg mL^-1^ ciprofloxacin was added to the first column of a 96 well plate and serially diluted 2-fold across the plate in MHB. The exponential phase cultures were diluted to 10^6^ colony forming units (c.f.u) mL^-1^ and 100 μL was added to each well to give a final *E. coli* concentration of 5 × 10^5^ c.f.u mL^-1^ and a final ciprofloxacin concentration of 1 μg mL^-1^ in the first column, down to 0.0019 μg mL^-1^ in the tenth column. Column eleven was a positive control of *E. coli* in MHB without ciprofloxacin and column twelve was a negative control of 200 μL MHB without *E. coli.* 96 well plates were then incubated for 24 hours at 37 °C on a shaking incubator at 200 rpm. The minimum inhibitory concentration (MIC) was determined using a plate reader (CLARIOstar®, BMG Labtech) and defined to be the concentration of ciprofloxacin that resulted in no growth at OD_600_ after blank correction.

Similar MIC tests were performed while intermittently maintaining the ciprofloxacin exposure. These tests were carried out by either adding ciprofloxacin to the overnight culture or not, and then adding it to the 1:40 dilution sub-culture or not. We therefore tested four different experimental conditions:

1. Adding ciprofloxacin to the overnight culture and to the sub-culture (Continuous exposure)
2. Adding ciprofloxacin to the overnight culture but not to the sub-culture (2h break)
3. Not adding ciprofloxacin to the overnight culture but to the sub-culture (17h break)
4. Adding ciprofloxacin neither to the overnight culture nor to the sub-culture (19h Break)

Cross resistance MIC tests were performed as above on a range of other antibiotics, adjusting antibiotic concentrations as appropriate. From the fluoroquinolone class we tested: finafloxacin, levofloxacin, moxifloxacin, and ofloxacin. From other antibiotic classes we tested: ampicillin (beta-lactam), tetracycline (tetracycline), trimethoprim (aminoglycoside), gentamicin (antifolates), and polymyxin B (polymyxins).

### Time-kill assays

Overnight cultures were prepared as described above, maintaining ciprofloxacin selection pressure as appropriate. Each culture was diluted 1:40 in LB and grown for 2 hours to reach exponential phase, again maintaining the same ciprofloxacin selection pressure. Once the culture had reached exponential phase, it was split to be challenged by 4 ciprofloxacin concentrations, i.e. 1×, 10×, 25×, and 100× the MIC_PS_ value for the structured environment mutant and 1×, 25×, 100×, and 200× WT the MIC_PS_ value for the well-mixed environment mutant. Cultures were incubated for 24 hours on a shaking incubator at 200 rpm at 37 °C and sampled in triplicate every hour. Samples were diluted in PBS and 10 μL spotted on LB agar plates for determination of colony forming units after overnight incubation of each plate at 37 °C. Means and standard error of measurements carried out in biological triplicate were calculated and graphs plotted using GraphPad Prism 9.

### Whole genome sequencing

Aliquots of the parental strain and resistant *E. coli* mutants were streaked from cryostocks on LB agar plates and grown overnight. A single colony was added to 100 μL of 1 × PBS and streaked out onto a second agar plate, thus covering 1/3 of the plate with a bacterial lawn that was incubated overnight at 37 °C. This lawn was transferred into a barcoded bead tube and sent to MicrobesNG for whole genome sequencing as previously described (Kraus et al. 2023). In brief, aliquots were taken from each tube and lysed by using 120 μL of TE buffer containing lysozyme (MPBio, USA), metapolyzyme (Sigma Aldrich, USA) and RNase A (ITW Reagents, Spain), and incubating for 25 min at 37 °C. Next, proteinase K (VWR Chemicals, Ohio, USA) (final concentration 0.1mg/mL) and SDS (Sigma-Aldrich, Missouri, USA) (final concentration 0.5% v/v) were added and incubated for 5 min at 65 °C. Genomic DNA was purified using an equal volume of SPRI beads and resuspended in EB buffer (10mM Tris-HCl, pH 8.0). DNA extracted was then quantified with the Quant-iT dsDNA HS (ThermoFisher Scientific) assay in an Eppendorf AF2200 plate reader (Eppendorf UK Ltd, United Kingdom) and diluted as appropriate. Genomic DNA libraries were prepared using the Nextera XT Library Prep Kit (Illumina, San Diego, USA) following the manufacturer’s protocol with the following modifications: input DNA was increased 2-fold, and PCR elongation time was increased to 45 seconds. DNA quantification and library preparation were carried out on a Hamilton Microlab STAR automated liquid handling system (Hamilton Bonaduz AG, Switzerland). Libraries were sequenced on an Illumina NovaSeq 6000 (Illumina, San Diego, USA) using a 250 bp paired end protocol. Reads were adapter trimmed using Trimmomatic version 0.30 with a sliding window quality cut-off of Q15. De novo assembly was performed on samples using SPAdes version 3.7, and contigs were annotated using Prokka 1.11.

The resultant reads were filtered and quality scores assessed with FastQC. Since the mean Phred scores were all > 30, no further filtering was applied. To identify mutations against the *E. coli* BW25113 reference genome (NCBI reference CP009273) we used the BRESEQ pipeline, with default settings in polymorphism mode (Deatherage and Barrick, 2014). BRESEQ uses Bowtie2 (Langmead and Salzberg, 2012) to map reads to a reference genome then recalibrates base quality scores using base position in the read. Single nucleotide polymorphisms (SNPs) were determined using a negative binomial model to determine SNP likelihood based on read-depth coverage. The default threshold frequency of 0.05 was used to identify variants. Deletions were determined from missing coverage and insertions and duplications from read junction evidence. Control samples were compared to the *E. coli* BW25113 reference genome to identify any pre-existing mutations and the reference updated before then comparing the resistant mutant strains to the updated reference to identify any mutations.

### Fabrication of microfluidic devices

Microfluidic mother machine devices were fabricated in polydimethylsiloxane (PDMS, Sylgard 184 silicone elastomer kit, Dow Corning) following previously reported protocols (Pagliara et al. 2011). Briefly, a 10:1 (base:curing agent) PDMS mixture was cast on an epoxy mold of the mother machine device kindly provided by S. Jun (Wang et al. 2010). Each mother machine device contains approximately 6000 lateral microfluidic channels with a width and height of 1 μm and a length of 25 μm. These channels are connected to a main microfluidic chamber that is 25 μm and 100 μm in height and width, respectively. After degassing, the PDMS mixture was allowed to cure at 70 °C for 2 h. The cured PDMS was peeled from the epoxy mould and fluidic accesses created using a 0.75 mm biopsy punch (RapidCore 0.75, Well-Tech) (Dettmer et al. 2014). After ensuring the fluidic accesses and the surface of PDMS chip were completely clean using ethanol wash, nitrogen gas drying and removal of any small particles using adhesive tape (Scotch® Magic™ Tape, 3 M), the PDMS chip, along with a glass coverslip (Borosilicate Glass No.1, Fisherbrand), were irreversibly sealed together as previously described (Łapińska et al. 2023). Briefly, both surfaces were exposed to oxygen plasma treatment (10 s exposure to 30W plasma power; Plasma etcher, Diener, Royal Oak, MI, USA), temporarily rendering the PDMS chip and glass coverslip hydrophilic. They were then immediately brought into contact with each other to bond them together. Next, the chip was filled with 5 μL of 50 mg mL^-1^ bovine serum albumin and incubated at 37 °C for 30 min, thereby passivating the internal surfaces of the device to preventing subsequent cell adhesion. A step by step protocol for the fabrication and handling of microfluidic devices can be found in (Cama and Pagliara 2021).

### Microfluidics-based time-lapse microscopy

Overnight cultures of either the 4-fold resistant triple mutant with mutations in the genes *acrR*, *marR* and *yhiJ* from the structured environment or the 16-fold resistant *gyrA* S83L mutant from the well-mixed environment were prepared as described above, maintaining ciprofloxacin selection pressure. A 50 mL aliquot from the overnight culture was then prepared for microfluidics experiments as previously described (Glover et al. 2022). Briefly, the aliquot was centrifuged for 5 min at 4000 rpm and 20 °C, the supernatant removed and filtered twice (Medical Millex-GS Filter, 0.22 μm, Millipore Corp.) to remove cellular debris and used to re-suspend the bacteria in their spent LB to an OD_600_ of 75. The suspension was then injected into the microfluidic mother machine device and incubated at 37 °C, the high bacterial concentration favouring the bacteria entering into the narrow side lateral channels from the main chamber (Bamford et al. 2017). An incubation time of around 5 – 20 minutes allowed, typically, between one and three bacteria to enter the side channels. A relatively shorter incubation time was needed for the more motile triple mutant from the structured environment. The microfluidic device set up was completed with the insertion of fluorinated ethylene propylene tubing (1/32” x 0.008”) into the fluidic accesses to connect the device to the inlet and outlet reservoirs which were attached to a computerised pressure control system (MFCS-4C, Fluigent) running MAESFLO software (Fluigent) as previously described (Cama et al. 2016). Once the incubation period was complete, spent LB was flushed through the microfluidic device at 300 μL h^-1^ for 8 minutes to remove any bacteria remaining in the main chamber and washed them into the outlet reservoir. The flow was then reduced to 100 μL h^-1^ for 2 hours for the initial growth phase on fresh LB. During this time, imaging of bacteria took place as the chip was mounted on an inverted microscope (IX73 Olympus, Tokyo, Japan) with automated stages (M-545.USC and P-545.3C7, Physik Instrumente, Karlsruhe, Germany) for coarse and fine movements, with a 60×, 1.2 N.A. objective (UPLSAPO60XW, Olympus) and a sCMOS camera (Zyla 4.2, Andor, Belfast, UK) (Goode et al. 2024). An area of interest of the camera was adjusted to visualise 23 side channels per image and images of 15 different areas of the microfluidic device were obtained at 30 minute time points. After the initial 2 hour growth phase, the antibiotic challenge phase began. This was initiated by switching the inlet reservoir from fresh LB to LB containing ciprofloxacin antibiotic at a concentration of 25× of the MIC_PS_ value (0.0375 μg mL^-1^), flowed through at 300 μL h^-1^ for 8 minutes, then 100 μL h^-1^ thereafter for 4 hours, continuing to take images every 30 minutes. After this, the regrowth phase began by removing the antibiotic and flushing fresh LB through the microfluidic device at 300 μL h^-1^ for 8 minutes before the rate was reduced to 100 μL h^-1^ for 4 hours. Imaging took place every 30 minutes to monitor regrowth for the structured environment mutant. Images were taken at the increased frequency of every 10 minutes for the well-mixed liquid mutant due to faster regrowth when switched back to fresh LB. The rate was then reduced to 50 μL h^-1^ and the setup left overnight before returning the next morning to take a final set of images.

Image analysis was performed in ImageJ software to calculate elongation rate and doubling time as previously reported (Lapinska et al. 2019). Briefly, elongation rate for each individual bacterium was calculated at each time point by first measuring the length of a bacterium and subtracting the length measured from the previous image to give the difference in lengths. This difference was then divided by the number of minutes elapsed between the two images to give an elongation rate in μm minutes^-1^. Wherever dividing bacteria were encountered, for the first frame of separation, the length of the parent cell was subtracted from the sum of the two daughter cells to give the length difference and then treated as separate cells from the subsequent frame onwards. When bacteria divided, doubling time was calculated by subtracting the time the parent bacterium was born from the time the bacteria split to form two daughter cells (Smith et al. 2019). All data were then analysed and plotted using GraphPad Prism 9.

### Statistical analysis

Normalised bacterial growth velocities for each environment were calculated by dividing each measured velocity for each exposure condition by the control velocity at every time point. Statistical significance was tested using unpaired, two-tailed, Welch’s t-test and standard deviation calculations were performed using GraphPad Prism 9. Error bars displayed in all graphs represent the standard error of the mean (SEM). Biological triplicate experiments were conducted in all instances, with sample sizes as defined in figure legends.

## Data availability

All data generated or analysed during this study are included in this published article and its supplementary information files.

## Author contributions

Conceptualization, S.P.; methodology, A.C., R.C. and S.P.; formal analysis, A.C., R.C., K.T.A. and S.P.; generation of figures, A.C. and S.P.; investigation, A.C., R.C., K.T.A. and S.P.; resources, S.P.; data curation, A.C., R.C., K.T.A. and S.P., writing – original draft, A.C. and S.P.; writing – review & editing, A.C., R.C., K.T.A. and S.P.; visualization, A.C and S.P.; supervision, R.C., K.T.A. and S.P.; project administration, S.P.; funding acquisition, S.P..

