## Supporting information for "Antibiotic resistant bacteria survive treatment by doubling while shrinking"

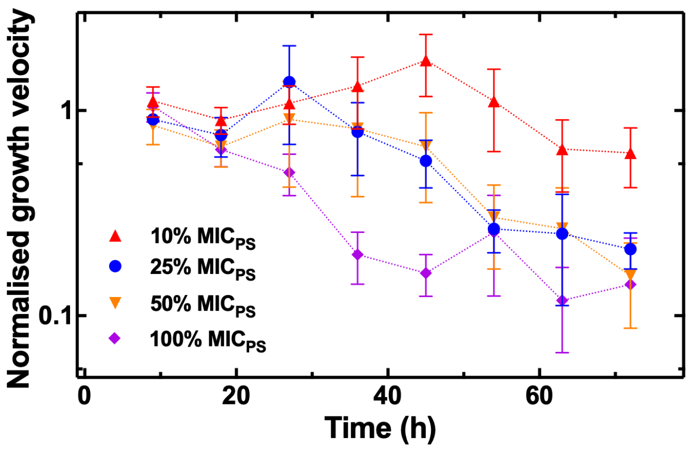


**Figure S1. Impact of the exposure to sub-MIC ciprofloxacin concentrations on bacterial growth in the structured environment.** Temporal dependence of *E. coli* growth velocity in the structured environment in the presence of different concentrations of ciprofloxacin: 10% (red upwards triangles), 25% (blue circles), 50% (orange downwards triangles), 100% (purple diamonds) the MIC of ciprofloxacin against the parental strain *E. coli* BW25113. Bacterial growth velocity values at each time point and for each condition were normalised to the bacterial growth velocity value measured for bacteria growing in the absence of ciprofloxacin at each time point. Symbols and error bars are the mean and standard error obtained by averaging measurements performed in biological triplicate.

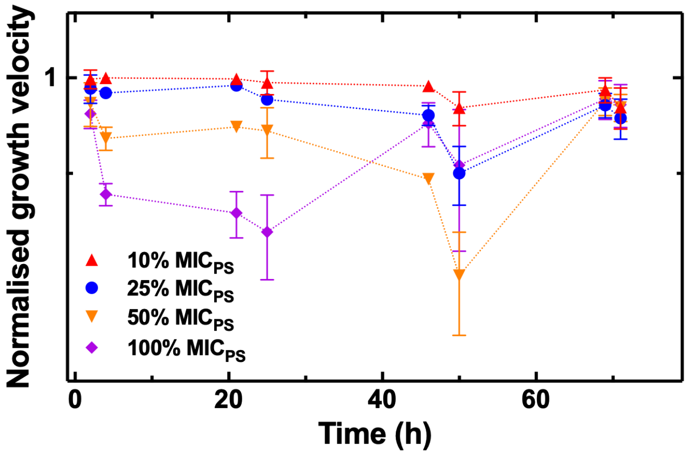


**Figure S2. Impact of the exposure to sub-MIC ciprofloxacin concentrations on bacterial growth in the well-mixed environment.** Temporal dependence of *E. coli* growth velocity in the well-mixed environment in the presence of different concentrations of ciprofloxacin: 10% (red upwards triangles), 25% (blue circles), 50% (orange downwards triangles), 100% (purple diamonds) the MIC of ciprofloxacin against the parental strain *E. coli* BW25113. Bacterial growth velocity values at each time point and for each condition were normalised to the bacterial growth velocity value measured for bacteria growing in the absence of ciprofloxacin at each time point. Symbols and error bars are the mean and standard error obtained by averaging measurements performed in biological triplicate.

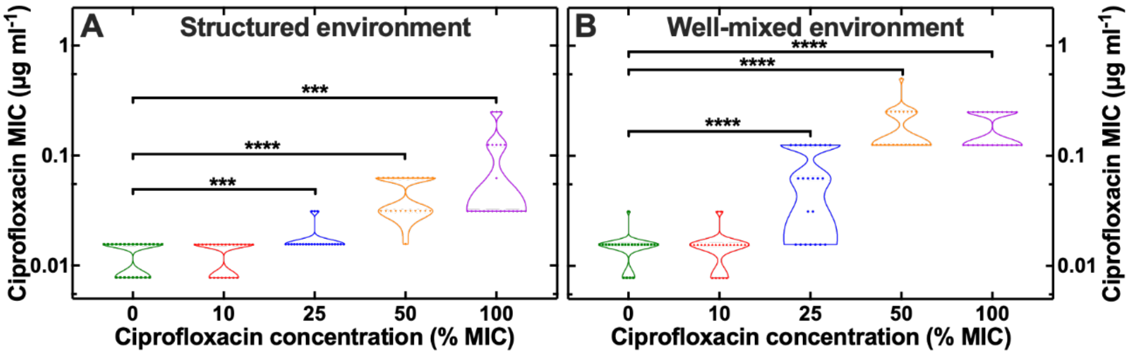


**Figure S3. Impact of the environmental structure on the emergence of genetic resistance to ciprofloxacin.** Dependence of the emergence of resistance to ciprofloxacin on the concentration of ciprofloxacin experienced by *E. coli* during evolutionary experiments in (a) the structured and (b) the well-mixed environment. Each symbol represents the ciprofloxacin MIC value measured for one out of eight technical replicates from three different evolutionary experiments for a total of 24 measured ciprofloxacin MIC values for each environmental condition. ***: $p < 0.001$; ****: $p < 0.0001$.

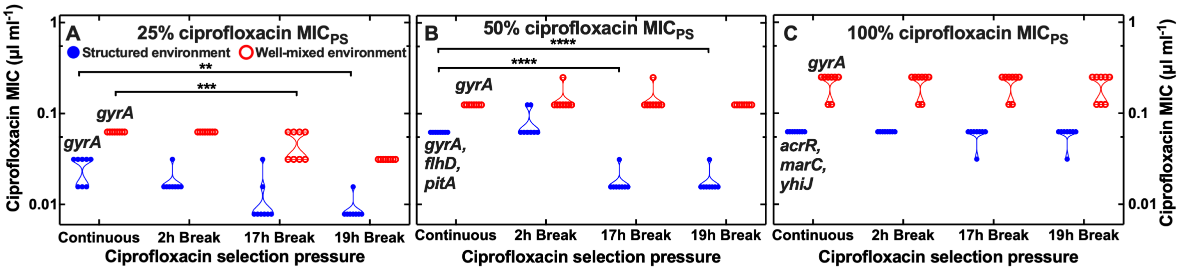


**Figure S4. Impact of the duration of ciprofloxacin selection pressure on the maintenance of resistance to ciprofloxacin.** Dependence of the capability of resistant mutants to maintain resistance to ciprofloxacin on the duration of ciprofloxacin selection pressure at the end of evolutionary experiments in the structured (blue filled circles) or well-mixed environment (open red circles) using ciprofloxacin at (a) 25%, (b) 50% or (c) 100% its MIC against the *E. coli* parental strain ($MIC_{PS}$). At the end of each evolutionary experiment, we either kept using ciprofloxacin (continuous) or removed ciprofloxacin from the environment either for 2 h (i.e. when the mutant culture was growing to exponential phase before the microbroth serial dilution assay), for 17 h (i.e. when the mutant culture was growing overnight for biomass expansion), or for 19 h (both during overnight culture and following exponential growth). Each symbol represents the ciprofloxacin MIC value measured in eight technical replicates for each mutant indicated in figure. **: $p < 0.01$; ***: $p < 0.001$; ****: $p < 0.0001$.

**Table S1. Molecular mechanisms underpinning the emergence of genetic resistance to ciprofloxacin.** Structure of the environment, experiment replicate number and ciprofloxacin concentration as a percentage of the $MIC_{PS}$ value employed during each triplicate evolutionary experiments. Corresponding MIC fold change of each mutant compared with the parental strain, mutation position, type, annotation, gene and description of gene product. The genome of each resistant mutant was sequenced and compared with the genome of *E. coli* BW25113 via our BreSeq pipeline. All mutations reported occurred with 100% frequency in the reads from sequencing. Mutations with frequency lower than 100% are reported in Table S2.

**Table S2. Further mutations occurring in each evolutionary experiment.** Structure of the environment and ciprofloxacin concentration as a percentage of the $MIC_{PS}$ value employed during each triplicate evolutionary experiments. Corresponding MIC fold change of each mutant compared with the parental strain, mutation position, type, frequency, annotation, gene and description of gene product.
